# Plant response to intermittent heat stress involves modulation of mRNA translation efficiency

**DOI:** 10.1101/2024.01.24.576986

**Authors:** Arnaud Dannfald, Marie-Christine Carpentier, Rémy Merret, Jean-Jacques Favory, Jean-Marc Deragon

## Abstract

Acquired thermotolerance (also known as priming) is the ability of cells or organisms to better survive an acute heat stress if it is preceded by a milder one. In plants, acquired thermotolerance has been studied mainly at the transcriptional level, including recent descriptions of sophisticated regulatory circuits that are essential for this learning capacity. In this work, we tested the involvement of polysome-related processes (translation and cotranslational mRNA decay (CTRD)) in plant thermotolerance using two heat stress regimes with and without a priming event. We found that priming is essential to restore the general translational potential of plants shortly after acute heat stress. We observed that mRNAs not involved in heat stress suffer from a reduction in translation efficiency at high temperature, whereas heat stress-related mRNAs are translated more efficiently under the same condition. We also show that the induction of the unfolded protein response (UPR) pathway in acute heat stress is favoured by a previous priming event and that, in the absence of priming, ER-translated mRNAs become preferential targets of CTRD. Finally, we present evidence that CTRD can specifically regulate more than a thousand genes during heat stress and should be considered as an independent gene regulatory mechanism.

## Introduction

Global climate change is a reality, day-to-day impacting our capacity to ensure food security worldwide (Duchenne-Moutien and Neetoo, 2021). Human activity is directly responsible for warming the world’s climate by 1.1°C compared to pre-industrial levels, and global warming is expected to reach 1.5°C by the early 2030s (IPCC, 2023). A direct consequence of these rises is an increase in the number of days per year with extremely high temperatures, a situation that threatens plant survival. Understanding how plants reprogram gene expression in response to heat is therefore crucial to developing the next generation of heat-stress resistant crops.

The reprogramming of gene expression involves many steps, from transcription and translation to post-translational modifications and degradation of RNA and proteins. To obtain an integrated view of the mechanisms regulating gene expression during heat stress, it is essential to obtain information on all these steps and the way they influence each other. Our current understanding of how plant gene expression is reprogrammed after a single heat stress event mainly concerns the gene transcription step (Kotak et al., 2007; Scharf et al., 2012; Jacob et al., 2017; Kerbler and Wigge, 2023). Briefly, upon heat stress, heat shock proteins (HSP)90/70 are redirected to unfolded proteins, allowing heat shock factor (HSF)A1 to enter the nucleus to displace histone variant H2A.Z and stimulate transcription of target genes (in cooperation with other transcription factors such as DRB2A and ABFs/AREBs). Induced proteins include secondary transcription factors (such as HSFA2, and HSFA3) that positively feedback on HSF and HSP transcription. An important function of HSP and other chaperone proteins is to restore protein homeostasis by preventing the accumulation of misfolded proteins in the cytosol via the cytosolic protein response (CPR) pathway (Sugio et al., 2009), and in the endoplasmic reticulum (ER), through the Unfolded Protein Response (UPR) pathway (Howell, 2021; Yu et al., 2022). The CPR and UPR pathways are also linked to protein degradation, preventing the precipitation of irremediably unfolded proteins (Chen et al., 2022; Krshnan et al., 2022; Lochli et al., 2022).

An important aspect of plant heat stress response is the ability of plants to withstand a severe heat stress if they have previously been exposed to a milder one (Baurle, 2016; Crisp et al., 2016; Shekhawat et al., 2022; Charng et al., 2023). This priming event allows plants to respond to a second acute event with a faster and stronger defense response, i.e. plants can ‘learn’ from the previous event. Plant stress memory (PSM) is defined as the ability of plants to maintain this primed state over time (Charng et al., 2023). The “memory phase” (i.e. the time between the first and second stress) can last from several hours to several days. Our understanding of the mechanisms leading to priming and PSM is mostly limited to the transcriptional and post-translational steps (Shekhawat et al., 2022; Charng et al., 2023). Following priming, HSFA1 can activate the transcription of genes that produce “effectors” (such as HSP101 and HSP21 (Harndahl et al., 1999; Queitsch et al., 2000)) that are responsible for maintaining the primed state in the memory phase and for modifying the outcome of the second acute stress. HSFA1 can also initiate the transcription of genes that produce secondary transcription factors (such as HSFA2 and HSFA3) and “regulators” that can either positively regulate effectors (maintainers, such as HSA32 (Wu et al., 2013)) or negatively (erasers, such as the protease FtsH6 (Sedaghatmehr et al., 2016)). Specific chromatin modifications allow the transcriptional re-induction of effector genes at the end of the memory phase (Lamke et al., 2016), while small RNAs are involved in the post-transcriptional regulation of heat-induced mRNAs throughout the heat stress period (Stief et al., 2014; Shekhawat et al., 2022). Therefore, sustained transcriptional activity driven by multiple integrated regulatory circuits and small RNAs are essential to establish PSM and acclimate plants. Recently, alternative splicing has also been shown to be involved in priming and PSM (Ling et al., 2018). Upon priming, splicing is repressed, leading to the accumulation of stress-responsive transcripts containing introns. Subsequent exposure to heat triggers the correct splicing of these transcripts and the production of the corresponding stress-responsive proteins (Ling et al., 2018; Ling et al., 2021).

Very little is known about the potential contribution of other post-transcriptional regulatory mechanisms (i.e. mRNA decay and translation) to priming and PSM. A recent report using yeast cells showed for the first time that both nuclear and cytoplasmic mRNA degradation contribute to gene expression memory (Li et al., 2023), but a similar role for mRNA decay in plants has not yet been explored. However, it has been reported that one of the fastest ways (within minutes) for plants to adapt their genetic program to rapidly fluctuating temperatures is to modulate global mRNA populations by inducing decay processes (Merret et al., 2013; Crisp et al., 2016). We have previously observed that a single brief heat stress can induce the partial decay of nearly 25% of the Arabidopsis transcriptome within a few minutes (Merret et al., 2013). This process, which is critical for plant survival, mainly targets “growth and development” functions and is likely to contribute to the establishment of the new incoming “heat stress” program. Another unexplored possibility is that heat stress simultaneously alters the ability of these destabilized mRNAs to produce corresponding proteins by reducing their translation efficiency. Thus, heat stress could not only reduce the global abundance of specific mRNAs by targeting them for degradation, but also reduce their capacity to produce corresponding proteins by limiting their translation efficiency. In contrast, the translation efficiency of pre-existing heat stress-responsive mRNAs could be favored under these conditions.

Recently, it has been shown that mRNAs associated with polysomes can undergo 5’-3’ exonucleolytic decay in a process called mRNA co-translational decay (CTRD) (Pelechano et al., 2015). This process, first discovered in yeast (Hu et al., 2009) but conserved in mammals and plants, affects all transcripts (Pelechano et al., 2015; Yu et al., 2016; Carpentier et al., 2020; Guo et al., 2023). It proceeds by decapping polysomal mRNAs, producing a 5’-monophosphate (5’P) end. The exonuclease XRN1/XRN4 then degrades the mRNAs as they are translated, following the last ribosomes. 5’P sequencing (5’P-seq) is an approach that allows the capture and sequencing of mRNA presenting a free 5’-phosphate end and is therefore used to measure CTRD intensity (Carpentier et al., 2021; Zhang and Pelechano, 2021). A 3 nucleotide periodicity is observed for 5’P-seq reads (also called degradome reads), reflecting the protection provided by elongating ribosomes. There is also a general accumulation of reads 16-17 nucleotides upstream of stop codons due to translation termination, which is slower than elongation. As 5’P-seq can reveal ribosome footprints *in vivo*, it can also be used to assess ribosome dynamics at the initiation, elongation, and termination steps (Nersisyan et al., 2020; Carpentier et al., 2021; Zhang and Pelechano, 2021). The discovery of the CTRD process also means that the translation efficiency of a given mRNA is best measured as the ratio of its amount in polysomes divided by the number of molecules actively degraded by CTRD, rather than by its amount in polysomes alone (Carpentier et al., 2020). However, it is not yet clear whether CTRDs represent a new regulatory mechanism that could regulate translational output independently of the total amount of mRNA in polysomes and whether this process could contribute to priming and PSM.

The aim of this work is to evaluate the importance of translation and CTRD in the regulation of plant gene expression under two heat stress regimes with or without priming. We found that priming preserves the global translational potential of the plant and differentially affects the translational efficiency of stress and non-stress related genes. We also found that priming is essential for up-regulating the UPR pathway and protecting ER-associated mRNA from CTRD. Finally, we provide evidence that CTRD intensity is not simply a consequence of more or less association with polysomes but acts independently during heat stress to regulate the expression of more than a thousand genes.

## Results

### Effect of priming on plant global translation potential

Variations in polysome gradient profiles are often used as a proxy to assess the overall translational potential at different stages of development or environmental conditions (Chasse et al., 2017). In a first attempt to measure the effect of priming on the global translational potential of plants, we used polysome gradient profiles to assess polysome levels at different time points of two heat stress regimes, with and without a priming event (Figure 1A). The Short Acquired Thermotolerance (SAT) and Basal Thermotolerance (BT) heat stress regimes have previously been used to study the physiological effects of priming (Yeh et al., 2012). For the SAT regime (Figure 1A), Arabidopsis plants were grown *in vitro* for 5 days at 20°C and collected 5 hours after daylight, just before exposure to the 1-hour 37°C priming event (p1). Plants were also collected after 15 min at 37°C (p2), after 2 h recovery at 20°C (p3), after 15 min of the 30 min 44°C heat stress period (p4), and after 7.5 h recovery at 20°C (p5). For the BT regime (Figure 1A), plants were collected before the 30 min 44°C heat stress period (pA), after 15 min at 44°C (pB) and after 7.5 h recovery at 20°C (pC).

**Figure 1:**
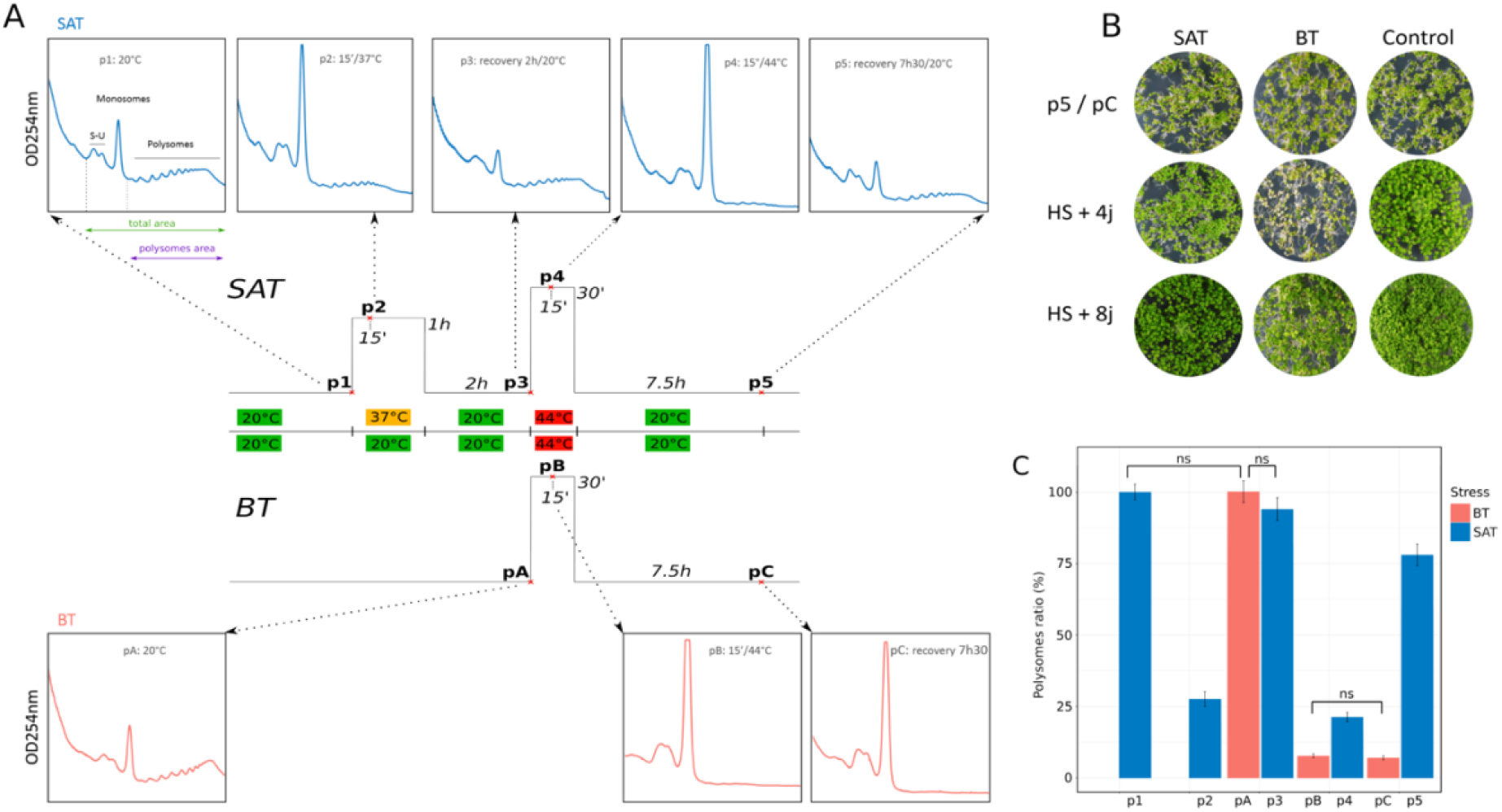
A 37°C priming step positively affects polysome levels at 44°C and during recovery. A) Scheme of SAT (Short-term Acquired Thermotolerance) and BT (Basal Thermotolerance) heat-stress regimes (Yeh et al., 2012). Plant samples were taken at different times during SAT (p1 to p5) and BT (pA to pC). (B) Phenotype of seedlings after SAT or BT treatment at time point p5 and pC, four and eight days after heat stress compared to control (same time points for plants always maintained at 20°C). C) Estimation of the proportion of polysomes at the different time points of SAT and BT compared to the 20°C situation (arbitrarily fixed at 100%). For each profile presented in A, the amount of polysomes was normalized to the total amount of ribosomes (i.e. 40S, 60S monosomes and polysomes) and both values were estimated by measuring the corresponding area above the baseline in our sucrose gradient. Data are the mean of three biological replicates. All variations are significant (Wilcoxon test, *p-value* < 0.05) except where indicated.

Plants at p5 and pC as well as control plants (i.e. always grown at 20°C) have the same global phenotype (Figure 1B and S1). However, as expected (Yeh et al., 2012), plants exposed to SAT have better long-term survival than plants exposed to BT (Figure 1B and S1). Four days after heat stress, cotyledons of plants exposed to BT are bleached, while plants exposed to SAT show only reduced growth compared to the control without any bleaching effects (Figure 1B and S1). Eight days after heat stress, most cotyledons of BT plants are dead, but a significant proportion of these plants can still recover (Figure 1B) and produce new leaves after ten days of recovery (Figure S1). These results indicate that Arabidopsis (Col0) can survive both heat stress regimes, but in contrasting ways.

Next, we generated polysome profiles using sucrose gradient sedimentation and OD_254nm_ measurement for each time point of SAT and BT and measured the proportion of polysomes at each time point compared to the 20°C condition (Figure 1A and C). As expected, heat stress induces a decrease in translational activity by dissociating polysomes from mRNA (Merret et al., 2015). Treatment at 37°C for 15 minutes (p2) induces a decrease in polysomes to about 25% of the amount present at 20°C. However, after two hours at 20°C (p3), plants can recover almost the total amount of polysomes present in the initial 20°C control condition as previously described (Merret et al., 2017)

Interestingly, when we compared polysomes present in plants exposed to 44°C for 15 min after priming at 37°C (p4, blue) or without priming (pB, red), we observed that polysome levels are significantly higher in the primed condition (Wilcoxon test, *p-value* < 0.05) (Figure 1C). This suggests that the 37°C priming step somehow protects the translational potential of the plant by limiting polysome dissociation at 44°C. However, the most striking effect is observed after 7.5 hours of recovery at 20°C. In the primed condition (p5, blue), polysomes are back to about 75% of the amount found in the 20°C control condition. However, this significant recovery is not observed in the non-primed condition (pC, red), as the polysome levels are still low and identical to the stressed 44°C condition (compare pB and pC). This result suggests that without priming, plants exposed to 44°C for 15 min cannot recover their translational potential after 7.5 h at 20°C and are probably still in a stressed state.

As described previously (Garre et al., 2018) we also used our 5’P-seq data (see Figure 3 and below) to monitor plant translation potential via SAT and BT (Figure 2). In the 5’P-seq meta-analysis, an increased amount of protected fragments upstream of the start site and a decreased amount of the 16-17nt peak upstream of the stop sites are diagnostic of reduced translational potential (Garre et al., 2018). This is what is observed at 37°C (p2) with a return to an almost normal situation after 2 hours of recovery at 20°C (p3) (Figure 2A). At 44°C, a strong reduction in translation is observed for both situations (p4 and pB), with a slightly stronger effect for pB (Figure 2B). After 7.5h at 20°C without priming, translation is still mainly down (pC), but is functional again at p5, although at a lower level than the initial 20°C situation (Figure 2C). These 5’P-seq results are consistent with our polysome profile quantification data, confirming that priming has a strong positive effect on the overall translational potential of plants in response to severe heat stress.

**Figure 2.**
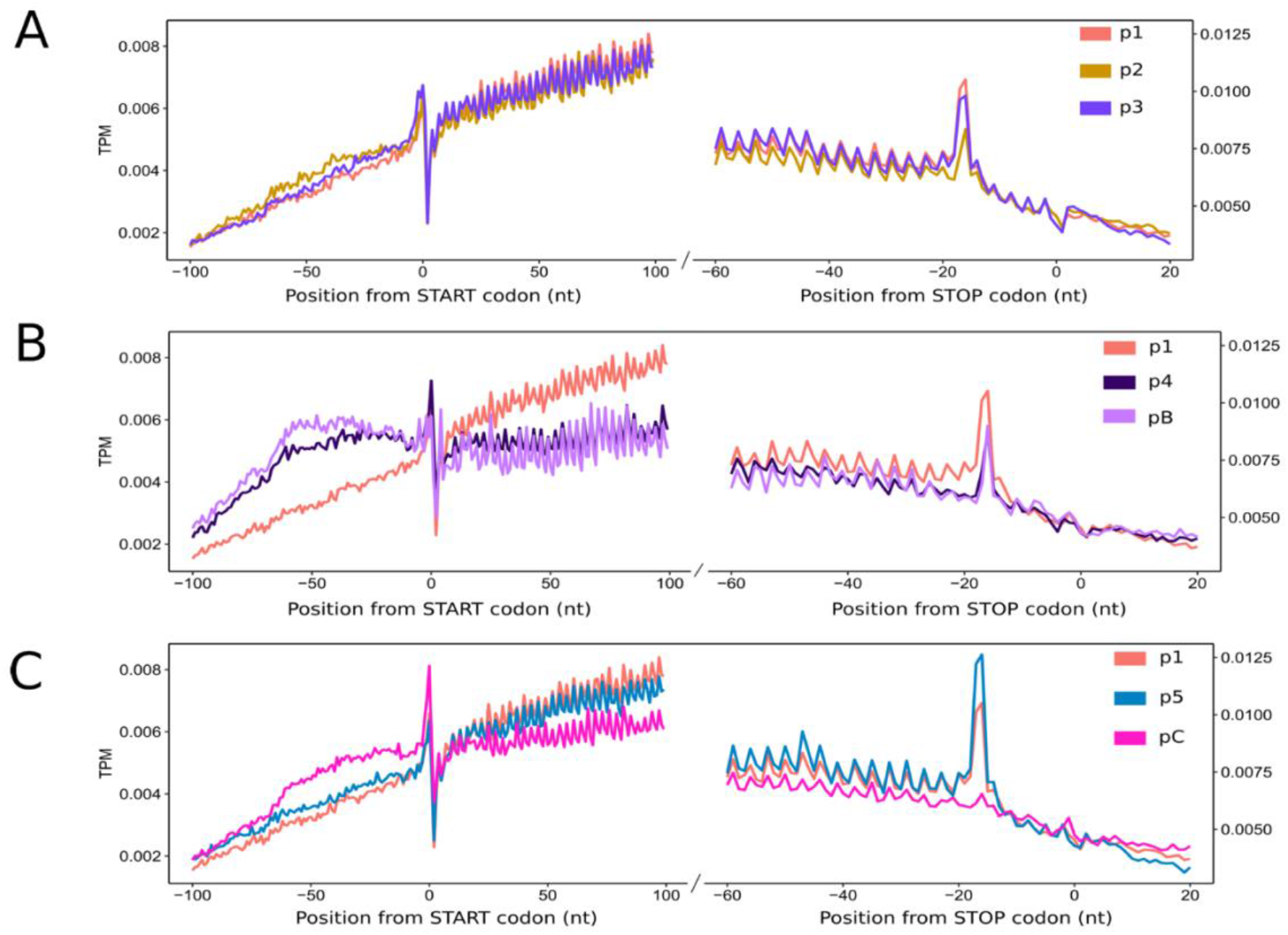
5’P-seq meta-analysis confirms the positive impact of priming on general translation levels. Meta-analysis of our 5’P-seq data. An increased amount of protected fragments upstream of the start site and a decreased amount of the 16-17nt peak upstream of the stop sites are diagnostic of a reduction in translation potential (Garre et al., 2018). A) This is what is observed at 37°C (p2) with a return to a near normal situation after 2h of recuperation at 20°C (p3). B) At 44°C a strong reduction of translation is observed for both situations (p4 and pB) with a slightly stronger effect for pB. C) Translation is still mainly down after 7.5h at 20°C without priming (pC) but is working again in p5 although at a lesser level than the 20°C initial situation. Globally the 5’P-seq results (A to C) confirm the polysome quantification data (Figure 1C).

**Figure 3:**
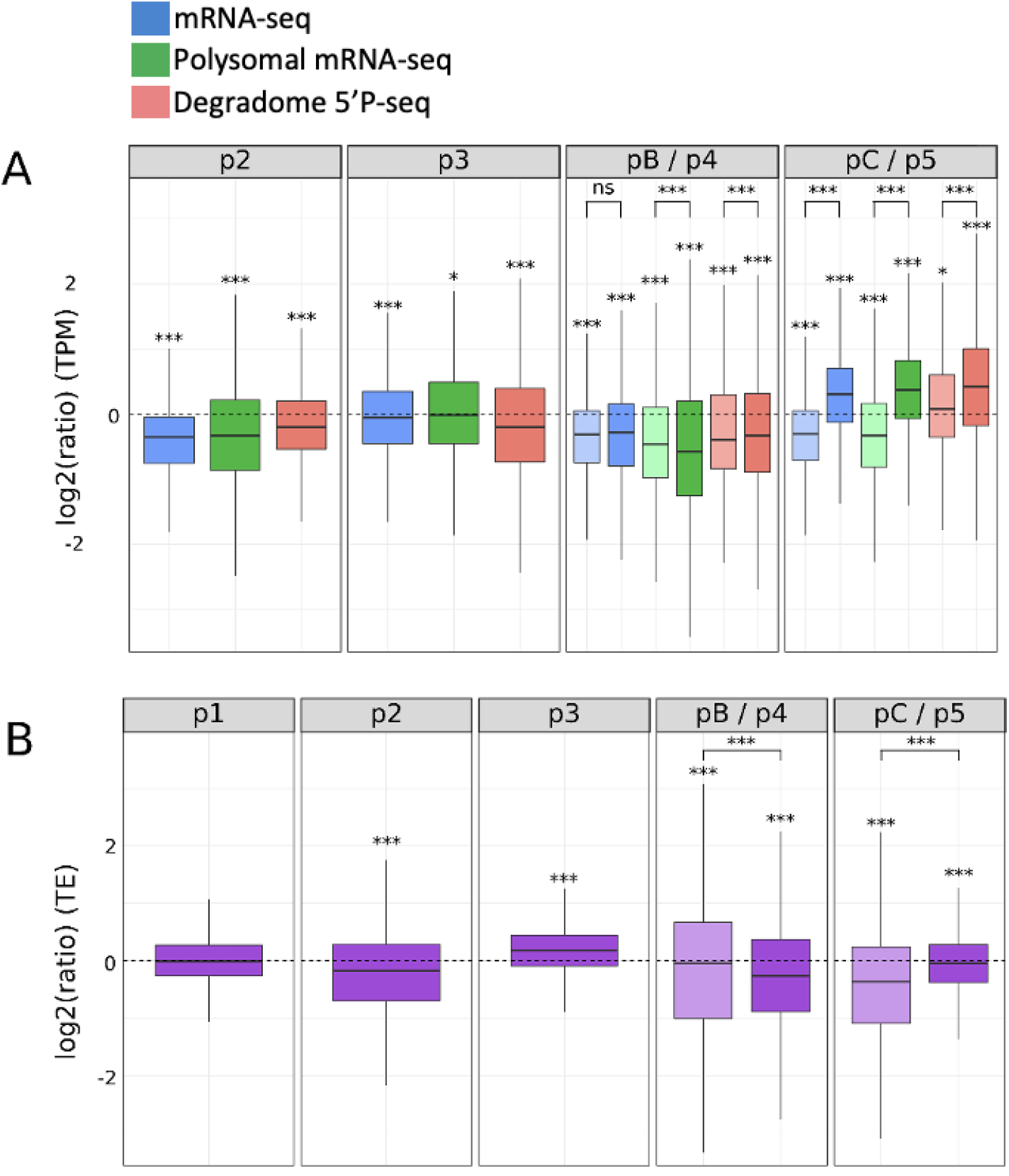
Variation in the distribution of total mRNAs, polysome-associated mRNAs, co-translational degradation products and translation efficiency levels at different time points of the SAT and BT heat stress regimes. A) Distribution (log2 (TPM ratio)) of a set of 13123 gene products present in at least 1 TPM (mean of three replicates) in all conditions (p1 to p5 and pA to pC) at different SAT and BT time points compared to the p1 situation (log2(TPM ratio) = 0). The distribution of total mRNAs is shown in blue, polysome-associated mRNAs in green and CTRD products in red. For p4/pB (44°C) and p5/pC (20°C recovery), light colored box plots represent unprimed conditions (pB and pC) and dark colored box plots represent primed conditions (p4 and p5). A Kolmogorov test was used to determine whether the distributions differ from 0 (stars above each distribution). A Wilcoxon test was used to compare primed and unprimed distributions (brackets above distribution pair) (* = p-value < 0.05, ** = p-value < 0.01, *** = p-value < 0.001). B) Variation in translation efficiency (polysomal mRNAs divided by degradation products) for the set of 13123 genes used in A at different SAT and BT time points compared to the p1 situation. A Wilcoxon test was used to compare P1 with the other distributions (stars above distributions) and to compare primed and non-primed distributions (brackets above distribution pair) (* = p-value < 0.05, ** = p-value < 0.01, *** = p-value < 0.001).

### Genome-wide analyses of the effect of priming on gene regulation: how important are polysome-related processes?

Next, we wanted to evaluate the importance of polysome-related processes (translation and CTRD) in the regulation of gene expression at different stages of SAT and BT heat stress. To this end, we performed a multiomics approach using a triple NGS strategy to analyze RNA samples collected at each SAT and BT time point (p1 to p5 and pA to pC) (see Figure S2 for a description of the experimental design and filtering strategy). We used an mRNA-seq approach to measure steady-state total mRNA levels, a polysomal mRNA-seq strategy to measure the global amount of mRNA associated with polysomes, and a degradome approach (i.e., 5’P-seq) to measure the amount of CTRD products (Figure 3A, in blue, green and red, respectively). A Principal Component Analysis (PCA) was used to evaluate the quality of replicates (Figure S3). The TPM values corresponding to the three replicates of all selected genes in the three databases are available in Supplementary Table 1.

While most studies use polysomal mRNA-seq to directly assess translation efficiency (TE), the ratio of polysomal mRNA-seq on degradome (5’P-seq) data provides a more realistic assessment of TE (Carpentier et al., 2020). Furthermore, the use of degradome data should allow us to determine whether CTRD represent an independent gene regulatory process or are merely a consequence of association with ribosomes.

In order to compare variations in the distribution of mRNAs produced by the same set of genes in SAT and BT, we decided to apply a filter to retain only those mRNAs that were present in at least 1 TPM (mean of replicates) in all conditions (p1 to p5 and pA to pC) (see Figures S2). Of these, 13123 genes were retained and used for further analysis (representing 50.3% of the total Arabidopsis genes number). GO analysis of these 13123 genes revealed that they are enriched in general functions involved in development and growth (i.e. major GO/Molecular Function terms: catalytic activity 10^E-104^, structural molecule activity 10^E-52^, structural component of ribosomes 10^E-34^; major GO/Biological Process terms: Metabolic process 10^E-112^, Cellular component organization 10^E-88^). This is to be expected in such a meta-analysis, as stress-responsive genes are in minority compared to other non-stress-responsive functions, and some stress-responsive genes will be filtered out of our gene set because they are not present at least at 1 TPM in all conditions.

Using these 13123 mRNAs, we performed a meta-analysis of their distribution compared to control distributions at 20°C (Figure 3A). This allowed us to measure variations in the distribution of total mRNAs, polysome-associated mRNAs and co-translational degradation products, thought to be SAT and BT. The specific effect of priming was also evaluated.

After 15 min exposure at 37°C (p2), there is a general tendency for a rapid decrease in the steady-state global mRNA level compared to the 20°C condition. As previously reported, this decrease is probably due to a general increase in global mRNA decay during heat stress, mainly targeting general functions involved in development and growth (Merret et al., 2013). We also observed a corresponding reduction in polysome-associated mRNAs and a smaller reduction in CTRD products, leading to a reduced TE level (Figure 3B, p1 vs. p2). This result shows for the first time that the previously observed decay at 37°C of a significant proportion of mRNAs (mainly involved in development and growth) is likely to correlate with a global reduction in their translational efficiency. Interestingly, after two hours of recovery at 20°C (p3), global and polysome-associated mRNA levels are almost back to their initial levels, while the rate of CTRD remains low. These results challenge the idea that CTRD is a passive mechanism that simply follows the rate of binding to polysomes and support the hypothesis that CTRD is an active/independent regulatory mechanism. Indeed, under these conditions, keeping the CTRD low while increasing the level of polysome-associated mRNA is a way to increase TE to even higher levels than in the initial 20°C situation (see p3 in Figure 3B, compare p1 to p3).

At 44°C, as at 37°C, a general reduction is observed for the three distributions in the unprimed (pB, light colors) and primed (p4, dark colors) conditions (Figure 3A). At this temperature, the strongest effect of priming is observed for polysome-associated mRNAs, whose level is lower compared to the unprimed condition (Wilcoxon test, *p-value* < 0.001), while the opposite is true for CTRD (Wilcoxon test, *p-value* < 0.001). Priming therefore has an effect by lowering the TE of our gene set at 44°C, a regulation that is less efficient without the priming step (Figure 3B, compare p1 to p4 and p1 to pB). After 7.5 hours of recovery at 20°C, the effect of priming (p5, dark colors) is very strong (Figure 3A). In the primed SAT regime, global and polysome-associated mRNA as well as CTRD are up, resulting in a TE almost back to the initial 20°C condition (p5 in Figure 3B). In the unprimed BT regime (pC, light colors), global and polysome-associated mRNA levels are still down (almost identical to the 44°C situation), while CTRD increases, leading to a decrease in TE (pC, Figure 3B). Again, polysomal mRNA levels and CTRD variations are anticorrelated, supporting the hypothesis that both processes could operate independently. We conclude that priming has a significant effect on gene expression, in part by decreasing the TE of our gene set (mainly non-stress related genes) at 44°C and increasing it during the recovery period.

### Effect of priming on heat stress-responsive genes

Another way in which priming could affect gene expression is by affecting the global composition of heat-inducible mRNAs that are up-regulated at 44°C. To test this, we first selected from our 13123 mRNAs in Figure 3 those that have a higher global steady-state level at 44°C (compared to the 20°C control condition) in a primed (N=730) or unprimed (N=197) regime. Surprisingly, a Venn diagram revealed that only 108 of these mRNAs are common to the primed and unprimed conditions (Figure 4A), demonstrating that the priming step has a very strong influence on the composition of heat-inducible mRNA populations following a 15-min exposure to 44°C. GO analysis of these common mRNAs identified “response to temperature stimulus” (GO: 0009266 P-value 7.28 10^E-38^) and “response to heat” (GO: 0009408, P-value 5.46 10^E-37^) as the most statistically supported GO terms. A closer look revealed that mRNAs of genes involved in the cytosolic protein response (CPR), a subcomponent of the general heat stress response (Sugio et al., 2009), were particularly enriched in this group (including HSP101, cytosolic HSP70s, HSP90s and HSP20s-like; see Supplementary Table 2). The CPR, as the unfolded protein response (UPR) occurring in the endoplasmic reticulum (ER), is induced by the accumulation of misfolded proteins in the cytosolic compartment and regulates protein folding and degradation, especially after stress.

**Figure 4:**
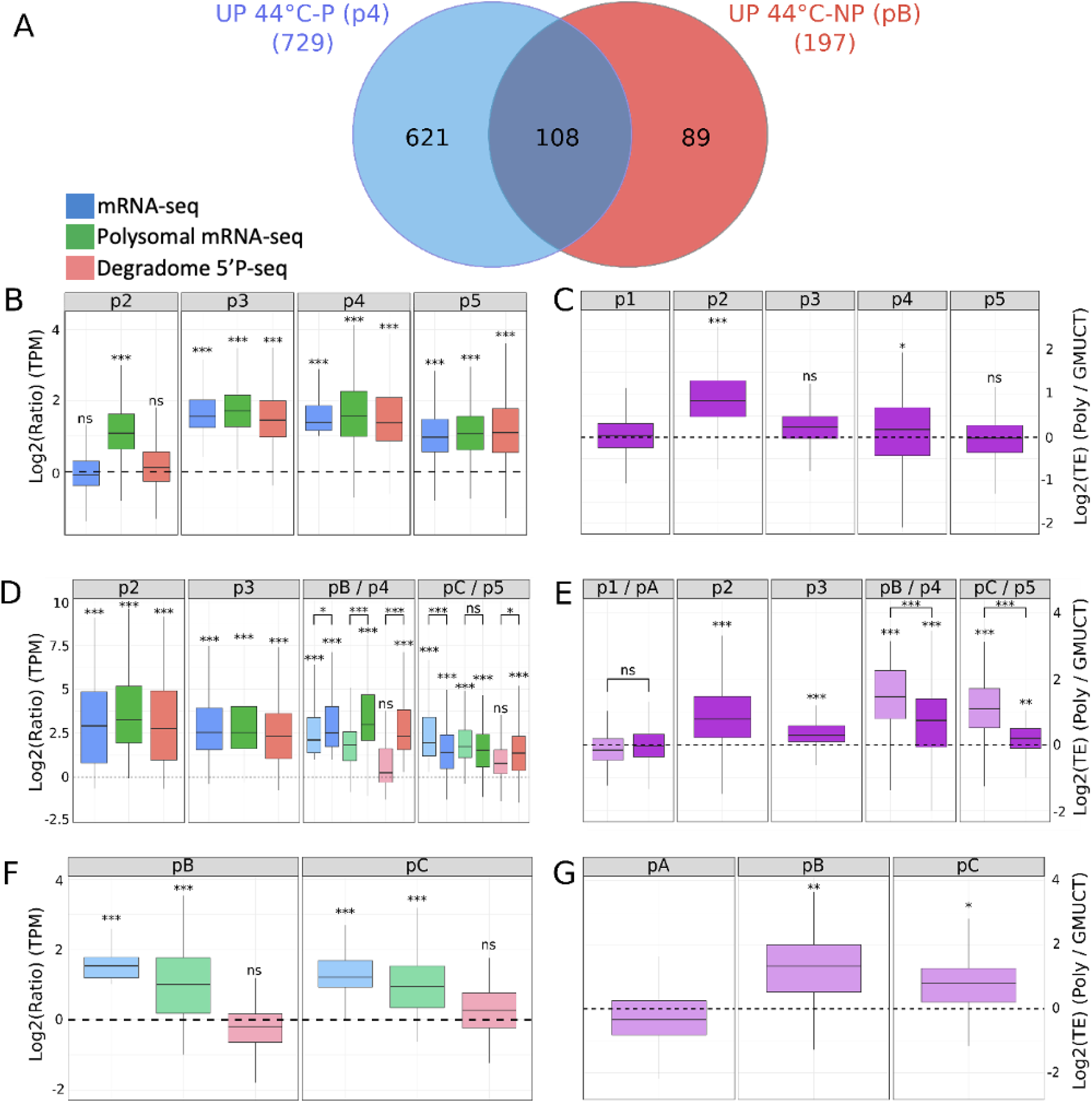
Effect of a 37°C priming step on mRNA populations and translation efficiency levels after 15 minutes of 44°C exposure. A) Venn diagram showing the number of mRNAs whose steady-state level is increased at 44°C after priming (p4), without priming (pB) and at the intersection of the two conditions. Only mRNAs with at least 1 TPM in all SAT and BT points were considered as starting material (see Figure S2). B) Distribution (log2(TPM ratio)) of a set of 621 mRNAs (only at 44°C after priming) at different SAT time points compared to the p1 situation (log2(TPM ratio) = 0). C) Boxplots showing the variation in translation efficiency (TE) of the 621 mRNAs during SAT (compared to p1). D) Distribution (log2(TPM ratio)) of a set of 108 mRNAs (up to 44°C with or without priming) at different SAT (dark colors) or BT (light colors) time points compared to the p1 situation (log2(TPM ratio) = 0). E) Box plots showing the variation of TE (compared to p1) of the 108 mRNAs during SAT and BT. F) Distribution (log2(TPM ratio)) of a set of 89 mRNAs (only up to 44°C without priming) at different BT time points compared to the p1 situation (log2(TPM ratio) = 0). G) Boxplots showing the variation in TE (compared to p1) of the 89 mRNAs throughout the BT regime. The distribution of total mRNAs is shown in blue, polysome-associated mRNAs in green and CTRD products in red. For B, D and F, A Kolmogorov test was used to determine whether the distributions differ from 0 (stars above each distribution), and a Wilcoxon test was used to compare primed and unprimed distributions (brackets above distribution pairs in D). For C, E and G, a Wilcoxon test was used to compare p1 with other distributions (stars above distributions) and to compare primed and unprimed distributions (brackets above distribution pairs in D). In all cases, p1 was used as the reference, as p1 and pA were compared and found to be equivalent (E). * = p-value < 0.05, ** = p-value < 0.01, *** = p-value < 0.001.

A similar GO analysis, this time on the 621 genes whose mRNAs specifically accumulate after priming, revealed that “protein folding” (GO:0006457, P-value 8.38 10^E-17^) and “response to endoplasmic reticulum stress” (GO:0034976, P-value 9.73 10^E-14^) are the most statistically supported GO terms. Major UPR players such as key regulators (AtbZIP60; ANAC089), key chaperones (BIP1 and 2, CALNEXIN 1 and 2, SHEPERD) and key members of the ERAD system (HRD1, DERLIN-1, CDC48 and UBPC6) are found in this group (Supplementary Table 2). This result suggests that 15 min of 44°C stress can induce the general and CPR heat stress response, at least at the global mRNA level, but that a previous 37°C priming event is required to additionally induce the UPR response.

GO analysis of the 89 genes whose mRNAs specifically accumulate without priming shows that “response to decreased oxygen levels” (GO:0036294, P-value 2.93 10^E-11^) and “response to stress” (GO:006950, P-value 1.14 10^E-10^) are among the most supported terms (Supplementary Table 2). Note that only two of these 89 genes can be directly associated with heat stress, suggesting that the 44°C stress, without priming, increases the range of stress-related genes induced.

We next decided to determine whether or not the observed increase in global mRNA levels at 44°C in SAT and BT was correlated with a corresponding increase in translational efficiency. For the 621 mRNAs whose accumulation is dependent on a priming event, the increase in steady-state mRNA levels at 44°C is associated with an increase in polysome-associated mRNAs and degradome fragments (p4, Figure 4B), leading to a small but significant (Wilcoxon test, *p-value* < 0.05) global increase in TE compared to the 20°C situation (Figure 4C, compare p1 to p4). This is in sharp contrast to the situation of non-stress-related genes, whose steady-state levels and translational efficiency are both downregulated at 44°C (Figure 3). We then focused on this subset of mRNAs throughout the SAT (Figure 4B). Surprisingly, we observed that in contrast to the 44°C situation (p4), these mRNAs do not show an increase in steady state mRNA levels at 37°C (p2). However, they show a strong increase in TE (Figure 4C, compare p1 to p2), which is mainly due to a strong increase in mRNA polysome association without a corresponding increase in co-translational degradation products (p2, Figure 4B). Therefore, the up-regulation of these genes at 37°C is only due to positive regulation at the translational level, a situation not detectable by simple transcriptomic analysis. This situation also illustrates again that CTRD is not the simple consequence of being associated with polysomes, since in this case the increase in polysomal mRNA does not lead to an increase in CTRD products. In the first recovery period (p3), all three distributions are up with a TE back to the initial 20°C situation (Figure 4B and C). The strong increase in steady-state mRNA levels observed in this first recovery period (p3) is likely due to positive feedback from at least some of the mRNAs that were massively involved in polysomes at 37°C (P2) (Figure 4B). In the final recovery period (7.5 h at 20°C, P5), all three distributions are still up, but TE is back to its initial level (p1), probably as the plants gradually switch from stress to a more balanced genetic program.

In contrast to the 621 mRNAs, the 108 mRNAs that globally accumulate to a higher level at 44°C in the SAT and BT regime (Figure 4D, pB/p4) do the same at 37°C (Figure 4D, p2). Furthermore, the TE level of these genes is increased at all stages of SAT and BT, with a higher increase in the unprimed (BT) compared to the primed (SAT) condition (Figure 4E). This higher increase in TE in the unprimed condition is mainly due to a lower level of CTRD degradation fragments compared to the primed condition at 44°C (pB/p4) and during the recovery period (pC/p5) (Figure 4D).

Finally, looking at the 89 mRNAs whose accumulation only occurs at 44°C in the BT regime (without priming), we observed an increase in TE (pB in Figure 4G), mainly due to a strong increase in the association of mRNAs in polysomes combined with the stability of the degradome fragments (pB in Figure 4F). During the recovery period (pC), the TE of these genes are still increased compared to the initial situation at 20°C (Figure 4G).

We conclude that the increase in global mRNA levels observed at 44°C in either SAT or BT is associated with an increase in TE, which is likely to lead to an increase in the production of corresponding heat stress responsive proteins.

### Co-translational decay regulates different subsets of transcripts through heat stress and recovery

Since we observed that CTRD does not necessarily correlate with mRNA levels in polysomes (Figures 3 and 4), we wondered to what extent CTRD could be used as a major mechanism to regulate gene expression in our two heat stress regimes. To answer this question, we applied a new filter to our initial mRNA set (N=13123), retaining only those mRNAs whose steady-state levels do not vary significantly during SAT and BT. This dataset (N=7929) was further subdivided into a new dataset, which we named CTRD-DB. For the CTRD-DB, we removed all transcripts that were differentially regulated in the polysomal mRNA database in at least one point of SAT and BT (i.e., 3929 transcripts). From the remaining 4000 transcripts, we kept only those whose abundance in the degradome database was significantly altered (compared to the 20°C control) in at least one SAT or BT step (p2 to p5 and/or pB to pC). We considered these selected 1126 transcripts to be the main targets of the CTRD pathway, as their levels vary significantly only in the degradome database during SAT and BT (see Figure S2).

Using the CTRD DB, we generated a heat map to visualize how these mRNA clusters and their CTRD levels vary during SAT and BT (Figure 5A). We found three significant clusters that were differentially regulated by CTRD. Cluster 1 mRNAs (N=602) are mainly degraded by CTRD during the 7.5 h recovery period, more so in SAT (p5) compared to BT (pC), while they are mostly preserved from CTRD at 44°C (pB/p4). GO analysis revealed an enrichment in this cluster for genes whose end products are localized in chloroplasts (GO: 0009507, 8.64 10^E-54^) and involved in small molecule metabolism (GO: 0044281, 4.68 10^E-27^). Cluster 2 mRNAs (N=223) are preferentially targeted by CTRD at 37°C but not at 44°C. GO analysis again revealed an enrichment for genes whose end products are localized in chloroplasts (GO: 0009507, 1.24 10^E-49^) and are involved in the generation of precursor metabolites and energy (GO: 0006091, 5.62 10^E-19^).

**Figure 5:**
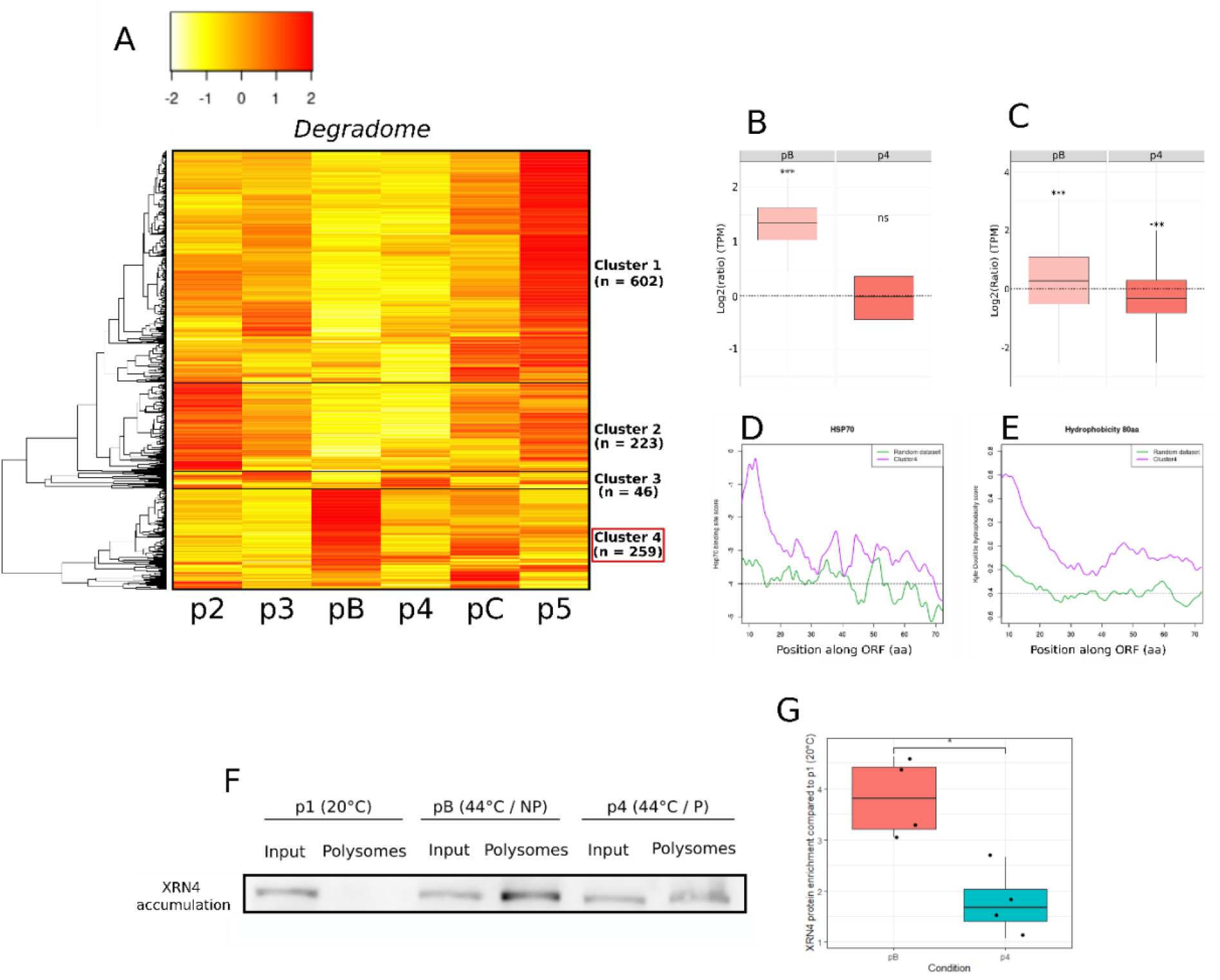
Co-translational decay regulates different subsets of transcripts through heat stress and recovery. A) Heat map of CTRD levels (n = 1126) at different stages of SAT and BT stress. Data are normalized to 20°C, red and yellow correspond to higher and lower CTRD levels, respectively, compared to the 20°C situation. B) Box plots showing the distribution of CTRD fragments of mRNAs from cluster 4 at 44°C in BT (pB) and SAT (p4). C) Boxplots showing the distribution of CTRD fragments of all mRNAs associated with the GO term “endoplasmic reticulum” (n = 944). D) Hydrophobicity scores of nascent peptides (10 to 70 aa) predicted for cluster 4 mRNAs and a control data set consisting of 259 randomly selected mRNAs. T-test was used to determine significant variations between two sets of genes (p-value = 2.64e-24) E) HSP70 binding score of nascent peptides (10 to 70 aa) predicted for cluster 4 mRNAs and a control data set composed of 259 randomly selected mRNAs. T-test was used to determine significant variation between two sets of genes (p-value = 3.19e-15). F) Western blot detection of XRN4 showing and enrichment in polysome fractions at 44°C in SAT and BT. G) Box plot showing the values for the 4 replicates of XRN4 enrichment in polysome fractions (Western blot). For each replicate, an equivalent amount of material was loaded for the input and polysome fractions and the quantity of XRNA in the polysome fraction was normalized by the signal detected in the input fraction. A Wilcoxon test was used to assess the significance of the difference between BT (pB) and SAT (pC) (*: p-value < 0.05).

Cluster 4 is particularly interesting as it shows a clear difference in CTRD targeting at 44°C between SAT and BT (Figure 5A). Indeed, cluster 4 contains 239 transcripts that are actively targeted by CTRD at 44°C in the BT regime (without priming) but not in the SAT regime (with priming) (compare cluster 4, pB and p4). GO analysis revealed that cluster 4 gene products are mainly localized in the endomembrane system (GO:0012505 endomembrane system, P-value 1.02 10^E-18^) and more specifically in the endoplasmic reticulum (GO:0005783, P-value 7.93 10^E-16^). A boxplot analysis of these 239 mRNAs confirms that they are degraded much more in BT than in SAT at 44°C (Figure 5B).

Previous work in human and mouse cell lines, as well as in Arabidopsis, has established a clear link between severe heat stress and early elongation ribosome pausing (when the nascent peptide emerges from the ribosomal exit tunnel), especially when this peptide is hydrophobic (Liu et al., 2013; Shalgi et al., 2013; Merret et al., 2015). This pause is likely due to the redistribution of HSP70 chaperones away from ribosomes during heat stress, as it can be alleviated by overexpressing HSP70 or exacerbated by HSP70 inhibitors (Liu et al., 2013; Shalgi et al., 2013; Merret et al., 2015). Hydrophobic nascent peptides are more dependent on HSP70s for proper folding and are therefore more affected by this pausing step. Since ribosome pausing is associated with the induction of CTRD and cluster 4 mRNAs are major targets of CTRD after severe heat stress in the absence of priming (Figure 5A), we decided to determine the hydrophobicity (Figure 5D) and HSP70 affinity scores (Figure 5E) of cluster 4 N-terminal peptides. We observed that cluster 4 genes encode highly hydrophobic nascent peptides with high HSP70 scores (Figure 5D, E). This result suggests that the high hydrophobicity of cluster 4 nascent peptides and their high requirement for HSP70s is probably the reason why they are preferential targets of CTRD at 44°C in the BT regime. But why does priming in the SAT regime prevent this? We have shown in this study that priming is necessary to induce the UPR response at 44°C, including the upregulation of ER-specific HSP70s (Bip1 and Bip2) (Figure 4, Supplementary Table 2). This increase in ER-specific HSP70s could limit ribosome pausing, explaining why CTRD is alleviated in the SAT regime for cluster 4 mRNAs. This conclusion is supported by Western blot analysis showing that XRN4, the major CTRD enzyme (Merret et al., 2015), localizes more efficiently to polysomes at 44°C in BT compared to SAT (Figure 5F and G).

Finally, we decided to see if this finding using cluster 4 genes could be generalized to all mRNAs in our initial dataset (N=13123) that are annotated as being translated in the RE (N=944). We observed that indeed all RE translated mRNAs are more efficiently targeted by CTRD in BT compared to SAT (Figure 5C). This suggests that although cluster 4 mRNAs are major targets of CTRD during BT, most mRNAs translated in the RE are likely to be at least partially regulated by CTRD in the same heat stress regime.

## Discussion

Thermotolerance (also known as priming) is a well-known phenomenon first discovered more than 40 years ago and defined as the ability of a mild heat shock treatment to protect cultured human cells from a more severe one (Gerner and Schneider, 1975). Mild heat shock has been shown in various animal cell culture systems to be important for the induction of HSP70 chaperones (Lindquist, 1981) and the stimulation of translation under severe heat shock conditions (Lindquist, 1980), two important features of thermotolerance. More recently, in mouse cells, pre-treatment with mild heat stress has been shown to limit the ribosome pausing effect induced by more severe heat stress (Shalgi et al., 2013). Thermotolerance is also well studied in plants, particularly the transcriptional regulation of heat shock proteins and their impact on plant survival (Martinez-Medina et al., 2016; Mauch-Mani et al., 2017; Charng et al., 2023). In this work, we used a multiomics approach with a three-pronged NGS strategy to investigate the role of polysome-related processes (translation and co-translational decay) in the acquisition of plant thermotolerance. First, using polysome gradient quantification and meta-analysis of degradome 5’P-seq data, we show that priming has a small positive effect on global translation in plants at 44°C, but a much stronger effect in the early recovery period (Figure 1 and 2). This priming-dependent spectacular rescue of plant translation potential only 7.5 h after severe heat stress was unexpected and is likely to be involved in the better long-term survival of plants exposed to SAT compared to BT. Indeed, the early increase in translation potential (almost to its 20°C level) is likely to be the key to restoring the growth and development program in SAT plants. This “solution” is not available to BT plants, which at the same time still have a very low translation potential (identical to the 44°C condition).

To further elucidate the importance of polysome-related processes in the regulation of gene expression, we generated transcriptome, translatome and degradome (5’P-seq) databases at all stages of the SAT and BT heat stress regimes. We started with a large dataset of genes that produce mRNAs that are present in at least 1 TPM at all SAT and BT steps (N=13123). This allowed us to monitor the regulatory behavior of ubiquitously expressed, mainly non-stress-related genes at the 5-day seedling development stage. As expected, we found that the steady-state level of these mRNAs decreases at 37°C, reflecting the decay process previously described under these conditions (Merret et al., 2013). We show here for the first time that this decrease in steady-state mRNA levels is associated with a significant decrease in translational efficiency (Figure 3). We suggest that down-regulation of these mRNAs at the translational level, rather than simply reducing their levels by decay, is important to switch from the “growth and development” to the “heat stress” program within minutes of exposure to 37°C. At 44°C, translation efficiency is again reduced, with a stronger effect for SAT than for BT, but the strongest effect of priming is in the recovery period, where translation efficiency can return almost to its 20°C level, a situation not observed for BT (Figure 3). In the SAT regime, the rescue of translation efficiency observed in our mRNA dataset, combined with the strong overall increase in plant translation potential at this stage (Figure 1 and 2), is likely to be important for restarting the “growth and development” program.

We next used the same strategy, but this time focusing on ubiquitously expressed genes whose steady-state level is significantly increased at 44°C in the SAT and/or BT. In contrast to the situation described above, these 44°C heat stress-inducible mRNAs (N=818) are not degraded at 37°C and show a clear increase in translation efficiency (and not a decrease as seen for non-stress-related genes). An increase in translation efficiency is also observed at 44°C, in combination with the observed increase in their global mRNA abundance. These results again point to a significant contribution of translation in positively regulating the onset of the heat stress response. However, the mechanism by which translation efficiency is increased is not equivalent for mRNAs up-regulated at 44°C only in the SAT regime (N=621) compared to those regulated only by BT (N=89) or by both regimes (N=108) (Figure 4A). Surprisingly, mRNAs up-regulated at 44°C only in the SAT regime are not up-regulated at 37°C in the mRNA-seq database (Figure 4B). Nevertheless, the translational efficiency of these mRNAs is strongly enhanced due to their increased presence in the polysome without a corresponding increase in the corresponding degradation fragments (p2 in Figure 4B). Thus, this population of mRNAs is only up-regulated at the translational level after a single 37°C exposure, a regulation that is not detectable by transcriptomic analysis. This 37°C translational up-regulation is likely to affect subsequent steps, as these mRNAs have increased steady-state levels in the first 20°C recovery period (p3) and through the SAT. We conclude from these analyses (Figures 3 and 4) that translation is an important regulatory step that contributes to the establishment of plant thermotolerance.

GO analysis of the 621 mRNAs that require priming for induction at 44°C revealed that they include a large proportion of genes involved in the UPR pathway (Howell, 2021) (Supplementary Table 2). This contrasts with the 108 mRNAs upregulated in both SAT and BT, which consist mainly of general heat stress genes and, more specifically, genes involved in the CPR pathway (Sugio et al., 2009) (Supplementary Table 2). This result suggests that the upregulation of the UPR in acute heat stress is dependent on a previous priming event, which is not the case for CPR. This is an unexpected consequence of priming that has not been previously documented and is likely to contribute to the thermotolerance property of the plant.

In this work, as previously recommended, we used a ratio between the translatome (polysome mRNA-seq) and degradome (5’P-seq) databases to better estimate translation efficiency (Carpentier et al., 2020). In addition, this allowed us to test whether the accumulation of CTRD fragments always correlated with the amount of mRNAs measured in polysomes. In many cases we observed a clear anticorrelation between the amount of mRNAs detected in polysomes and the accumulation of the corresponding CTRD fragments (the most obvious situations are Figure 3A, p2, p3, pC; Figure 4B, p2; Figure 4D, pB, pC; Figure 4F, pB, pC). These observations suggest that CTRD is not simply a neutral consequence of the presence of mRNAs in polysomes but could be used as an independent mechanism to regulate gene expression. In support of this proposition, we found that 1126 transcripts vary only in the degradome (5’P-seq) database in SAT and/or BT (Figure 5), suggesting that these mRNAs are exclusively regulated by CTRD in our experimental setting. Among these preferential targets of CTRD, we found a group of mRNAs (Figure 5, cluster 4) that are predominantly translated in the ER. These mRNAs are targeted by CTRD at 44°C in BT but not in SAT (Figures 5A and B). We propose that this difference between SAT and BT results from the inability to induce the UPR pathway in BT and to increase the amount of HSP70s (BIPs) chaperones in the ER (see above). Indeed, cluster 4 mRNAs encode proteins with highly hydrophobic nascent peptides that require HSP70s for folding (Figures 5D and E). In the SAT regime, the increased concentration of BIPs in the ER due to UPR induction helps to fold these hydrophobic nascent peptides, preventing ribosome pausing and the induction of CTRD for these messages (Liu et al., 2013; Shalgi et al., 2013; Merret et al., 2015). This cannot occur in BT because the UPR pathway is not stimulated in this condition, leading to a higher level of CTRD. We also observed that the situation described for cluster 4 mRNAs, which are likely to be regulated only by CTRD in SAT/BT (N=259), can be extended to all mRNAs translated in the RE (N= 944) (Figure 5C). In support of this model, we observed that XRN4, the major enzyme responsible for CTRD (Merret et al., 2015), is present at higher levels in the BT polysome at 44°C compared to SAT (Figure 5F and G).

In conclusion, we show here that polysome-related processes (translation and CTRD) are important to regulate gene expression after heat stress with or without a priming event. We observed that induction of the UPR in acute heat stress requires a previous priming event at a milder temperature and that in the absence of priming, ER-translated mRNAs become preferential targets of CTRD. Finally, we present evidence that CTRD should be considered as a novel gene regulatory mechanism that does not necessarily correlate with mRNA polysome association.

## Methods

### Plant material, growth conditions and heat stress

In all experiments, the wild-type Columbia ecotype was used. Plants were grown *in vitro* for 5 days on synthetic Murashige and Skoog (MS) medium (MS0213, Duchefa) containing 1% sucrose and 0.8% plant agar at 20°C under a 16h light (120 µmol.m-2.s-1)/8h dark cycle. Heat stress was applied in a water bath. For SAT, plates were transferred from the *in vitro* chamber to the water bath for 1 hour at 37°C, then transferred back to the chamber for 2 hours before returning to the water bath for 30 minutes at 44°C. Plates were then returned to the chamber for recovery. For BT, plants were transferred directly from the chamber to the water bath for 30 minutes at 44°C and then returned to the chamber for recovery. The different heat stresses were carried out in the dark and recovery was carried out under the same conditions as initial growth.

### Polysome gradients

Polysome profiling was performed as previously described (Merret et al., 2013). For polysome quantification, 100 mg ground powder (5-day-old seedlings) was incubated for 10 min on ice with 3 volumes of lysis buffer (200 mM Tris-HCl, pH: 9, 200 mM KCl, 25 mM EGTA, 35 mM MgCl2, 1% (v/v) detergent mixture [1% (v/v) Tween 20, 1% (v/v) Triton, 1% (w/v) Brij35 and 1% (v/v) Igepal], 1% (w/v) sodium deoxycholate, 0. 5% (w/v) polyoxyethylene tridecyl ether, 5 mM dithiothreitol, 50 μg mL-1 cycloheximide, 50 μg mL-1 chloramphenicol and 1% (v/v) protease inhibitor cocktail (Sigma-Aldrich)). The extract was centrifuged at 16,000g for 10 min, and 300 µL of the supernatant was loaded onto 9 mL 15-60% sucrose gradients. Ultracentrifugation was performed at 38,000 rpm (180,000g) for 3 hours using an SW41 rotor. Polysome profiles were recorded using an ISCO absorbance detector at 254nm and sucrose gradients were collected in 12 fractions of 650µL each: fractions 1-4 containing free mRNPs, fractions 5-8 containing ribosomal subunits and monosomes and fractions 9-12 containing polysomes. Absorbance at 254 nm was recorded using PeakTrack software and polysome profiles were plotted using Excel. The total amount of ribosomes was evaluated by measuring the area between the 40S, 60S, monosomes and polysomes profile and the basal lines, while the number of polysomes was evaluated by measuring the area on the polysome part of the profile only. The ratio of polysomes to total ribosomes at 20°C was arbitrarily set to 100%. Values represent the mean of three biological replicates.

For polysomal RNA extraction, 400 mg of ground powder (5-day seedlings) was used, following the same procedure as described above for purification of polysome fractions. The polysome fractions were pooled and SDS and EDTA were added to a final concentration of 0.1% and 4mM respectively. One volume of phenol/chloroform/isoamyl alcohol (25:24:1) was then added, the solution vortexed for 1 minute and centrifuged at 16,000g for 15 minutes. The phenolic phase was retained and one volume of chloroform/isoamyl alcohol (25:1) was added, the solution was vortexed for one minute and centrifuged at 16,000g for 15 minutes. The phenolic phase was retained, and 2 volumes of 100% ethanol were added, followed by centrifugation at 16,000g for 15 minutes. All centrifugations were performed at room temperature to avoid sucrose precipitation. The pellets were washed twice with 70% ethanol and resuspended in 30 µL of cold RNAse-free water.

### Total RNA extraction and NGS librairies

For total RNA extraction, 50 mg of ground powder (5 day seedlings) was used. One volume of phenol/chloroform/isoamyl alcohol (25:24:1) was added, the solution was vortexed for one minute and centrifuged at 16,000g for 15 minutes. The phenolic phase was retained and one volume of chloroform/isoamyl alcohol (25:1) was added, the solution was vortexed for one minute and centrifuged at 16,000g for 15 minutes. The phenolic phase was retained and 2 volumes of 100% ethanol were added followed by centrifugation at 16,000g for 15 minutes. The pellets were washed twice with 70% ethanol and resuspended in 50 µL of cold RNAse-free water.

All NGS experiments were performed in biological triplicates (i.e. different batches of plantlets collected and stressed at 1-week intervals), either on total RNA fractions or on polysomal RNA fractions (see above). All purified RNA extracts were DNAse treated using the Ambion Turbo DNAse Kit (Life technology). Quantities and quality of RNA were assessed using the Qubit Agilent 2100 Bioanalyzer and Plant RNA Nano Chip, respectively.

For RNA-Seq (total and polysomal), 1 µg of RNA was used for RNA library preparation using either total or polysomal RNA. Libraries were prepared using a NEBNext Poly(A) mRNA Magnetic Isolation Module and a NEBNext Ultra II Directional RNA Library Prep Kit for Illumina (New Enlgand Biolabs, #E7760S/L).

The degradome (5’P-seq) libraries were prepared as described previously (Carpentier et al., 2021). Briefly, 50 µg of total RNA was subjected to two rounds of poly(A+) purification. After ligation of 5ʹ adaptors, excess adaptors were removed by a new round of poly(A+) purification. Reverse transcription was performed using the SuperScript IV system according to the manufacturer’s instructions. cDNAs were amplified by 11 cycles of PCR. Libraries were purified using SPRIselect beads prior to quality control and normalization. Library quality was checked using an Agilent High Sensitivity DNA Kit (Agilent). Libraries were normalized, multiplexed and sequenced on NextSeq 550 (Illumina) in 75 pb single reads.

### Statistical analyses

All data presented were tested for normality using the Shapiro test. Depending on the result, a parametric (t-test) or non-parametric (Wilcoxon test) test was used to compare significance between conditions. For the quantification of polysome profiles (Figure 1C), t-tests were performed to test for significant differences between conditions. For Figure 3 and 4, two-sample Kolmogorov-Smirnov tests were used to compare distributions against 20°C and non-parametric Wilcoxon tests were used to compare distributions between primed and non-primed conditions (pB vs. p4 and pC vs. p5). All tests were performed using RStudio software.

### Bioinformatic analysis

Raw reads were trimmed using Trimmomatic v0.39 (Bolger et al., 2014). For RNA-Seq data, adapters and low quality reads were removed. For 5’P-seq data, reads were trimmed to 50nt. Trimmed reads were filtered from reads corresponding to chloroplast, mitochondrial, ribosomal and small RNA sequences using bowtie2 v2.4.4 (Langmead and Salzberg, 2012) in ‘sensitive-local’ mode. Remaining reads were mapped to the TAIR10 genome and corresponding gtf file annotations (Araport11) using Hisat2 v2.2.1 (Kim et al., 2019) with standard parameters. Only unique mapped reads were retained using samtools v1.13 with the ‘-q 10’ option (Li et al., 2009). Read counts were performed using htseq-count v1.99.2 in ‘union’ mode (Anders et al., 2015). Transcripts per million (TPM) values for each gene were calculated using the following methods: First, the read counts are divided by the length of each gene in kilobases (rpk), second, all rpk values in a sample are summed and divided by 1,000,000 (’per million’ scaling factor) and third, divide the rpk values by the per million scaling factor (TPM). For 5’P-seq data, unique alignment bam files are analyzed by the FivePSeq software (Nersisyan et al., 2020). Five TAIR10 chromosomes and Araport11 gtf are used as reference files.

A first filter was applied to the original dataset of 37390 genes: only transcripts containing ≥ 1 TPM in all studied conditions (p1 to p5 and pA to pC) were retained, resulting in the dataset of 13123 genes used in Figure 2 to 4. Quality of replicates was determine by a PCA analysis. Differential gene expression (DEG) analysis was performed on this dataset using DESeq2 (BiocManager) for statistical analysis (*p-value* < 0.05). Only DEG genes with TPM fold-change (FC) values > 2 at 44°C (compared to 20°C reference) were declared as significantly up-regulated and used in Figure 4.

To generate the CTRD-specific database of Figure 5, DEGs (up or down) in the RNASeqTotal database (5194 genes) or the RNASeqPolysomal database (3929 genes) were filtered out, resulting in a new dataset of 4000 genes. A further DEG analysis was then performed on these 4000 genes using the 5’P-seq database and 1126 genes were found to be differentially regulated, retained and considered to be major targets of CTRD. All GO analyses were performed using the Gprofiler website (Raudvere et al., 2019).

### Heat map, hydrophobicity and HSP70 scores

Hydrophobicity and HSC/HSP70 scoring were performed as described previously (Shalgi et al., 2013). Protein sequences encoded by cluster 4 (which are targets of XRN4 in the pB state) and a random population (of equal size; n = 258) were downloaded from TAIR10 and scores were calculated. Briefly, Kyte-Doolittle hydrophobicity scores and HSC/HSP70 binding scores were each calculated on 13 amino acid long windows shifted by one amino acid from the N-terminus to the C-terminus of the first 80 amino acids of each protein encoded by the two datasets examined (cluster 4 and random dataset). Mean values were calculated in each class from the values obtained for each gene, and the curves were fitted with the cubic spline interpolation mathematical function. A T-test was performed between each pair of populations.

### Western blot analyses

For polysomal-XRN4 western blots, proteins were extracted from sucrose fractions (1 to 12) after sucrose gradient sedimentation (see above) using 2 volumes of absolute ethanol. After 6hours of incubation at 4°C and centrifugation, protein pellets were washed and resuspended in Laemmli 4X buffer (0.25M Tris-HCl pH 6.8, 8% SDS (v/v), 40% glycerol (v/v) and 0.002% bromophenol blue (w/v), 5% beta-mercapto-ethanol (v/v)). After incubation at 95°C for 5 minutes. Proteins were separated on SDS-PAGE gels and electrotransferred to polyvinylidene fluoride membranes. Immunoblotting was performed in Tris-buffered saline (TBS), 5% skimmed milk (w/v), 1% Tween (v/v). XRN4 primary antibodies was diluted 1/5000 (Merret et al., 2013) and incubated overnight at 4°C with agitation. After incubation, the membranes were washed six times in TBS, 1% Tween (v/v). Secondary antibody (rabbit) was incubated in TBS, 5% skim milk (w/v), 1% (Tween) (v/v) for one hour at room temperature with agitation. The membranes were washed six times with TBS, 1% Tween (v/v). Development was performed using the Millipore Immobilon-P kit. Images were captured using the Fusion FX imaging system (Vilber). All blots were prepared and immunoblotted in parallel and simultaneously exposed for chemiluminescence quantification. For each of the 4 replicates, an equivalent amount of material was loaded for the input and polysome fractions and the quantity of XRNA in the polysome fraction was normalized by the signal detected in the input fraction.

#### Databases accession numbers

PRJNA1044736 (mRNA-seq), PRJNA1044743 (Polysome mRNA-seq) and PRJNA1044724 (Degradome (5’P-seq))

## Author Contributions

A.D., R.M., J.J.F. and J.M.D designed the research. A.D., R.M. and J.J.F. performed the experiments. A.D., M.C.C., R.M., J.J.F and J.M.D. analyzed the data. A.D., R.M., J.J.F and J.M.D. wrote the article.

## Acknowledgments and funding

This work was supported by a grant from the Agence National de la Recherche (ANR-22-CE20-0023-01), by a “New Frontier” grant from the Laboratoires d’Excellences (LABEX)” TULIP (ANR-10-LABX-41), by the CNRS and Université de Perpignan via Domitia. A.D. Salary was financed by the Occitanie Region. This study is set within the framework of the École Universitaire de Recherche TULIP-GS (ANR-18-EURE-0019)

