## Supplementary Figures for "Plant response to intermittent heat stress involves modulation of mRNA translation efficiency"

**Supplementary Figure 1**

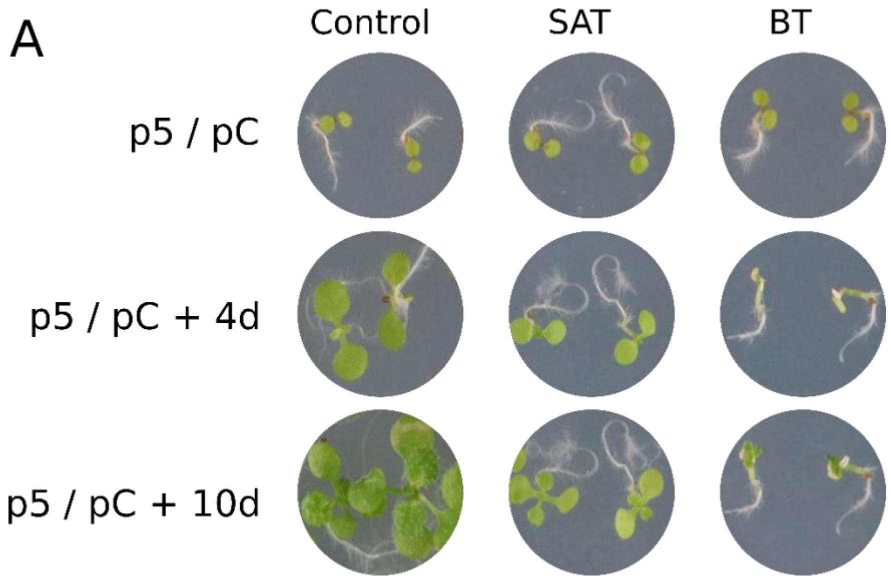


**Supplementary Figure 1**. Effect of priming on Arabidopsis plantlet phenotypes. For the SAT and BT heat stress regimes, images were taken 7.5 h (p5/pC of Figure 1), 4 days and 10 days after 44°C exposure. The control condition (left) is plants maintained at 20°C for the same time periods.

**Supplementary Figure 2**

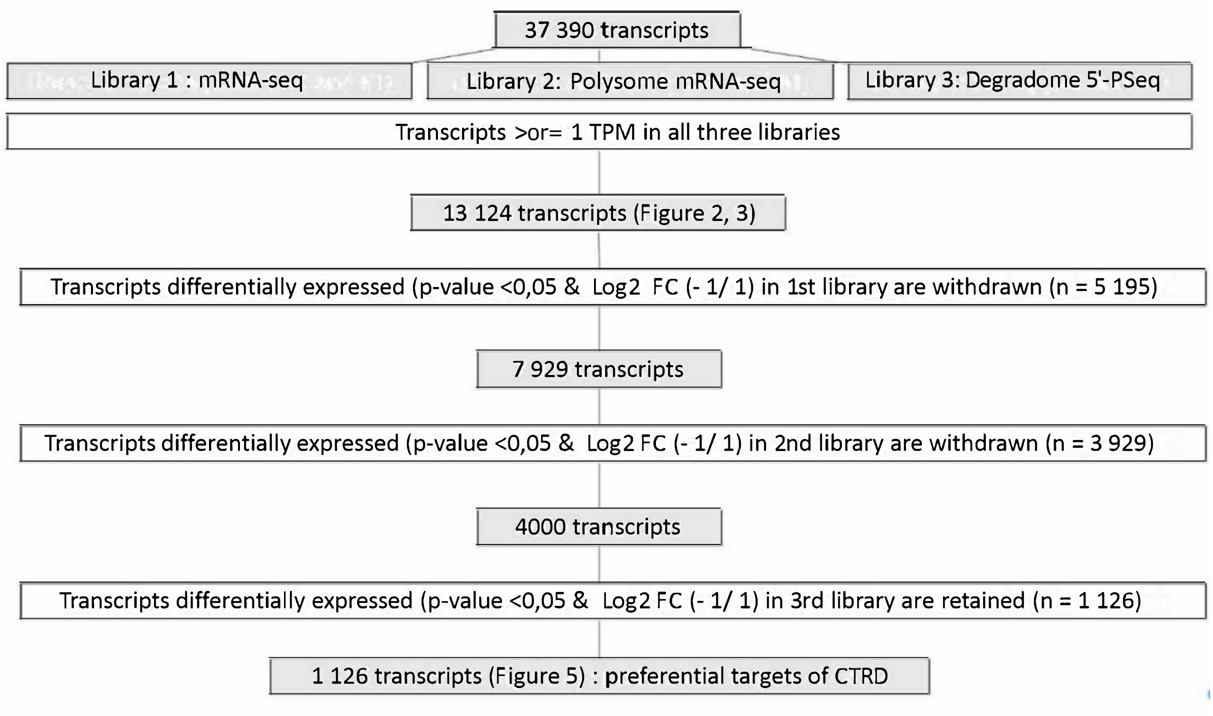


**Supplementary Figure 2.** Experimental design and filtering strategy used to generate Figures 2, 3 and 4. Three libraries were generated using different NGS techniques. A first filter was used to retain only transcripts with > 1TPM in all three libraries, generating a new dataset of 13,124 transcripts. The 5PSeq data was used to generate Figure 2 and the three datasets were used to generate Figure 3. To generate Figure 4, all transcripts differentially expressed (p-value <0.05 & FC -2/2) in libraries 1 and 2 were removed, resulting in a dataset of 4000 genes. Then, only transcripts differentially expressed in library 3 were retained and considered as preferred targets of CTRD.

**Supplementary Figure 3**

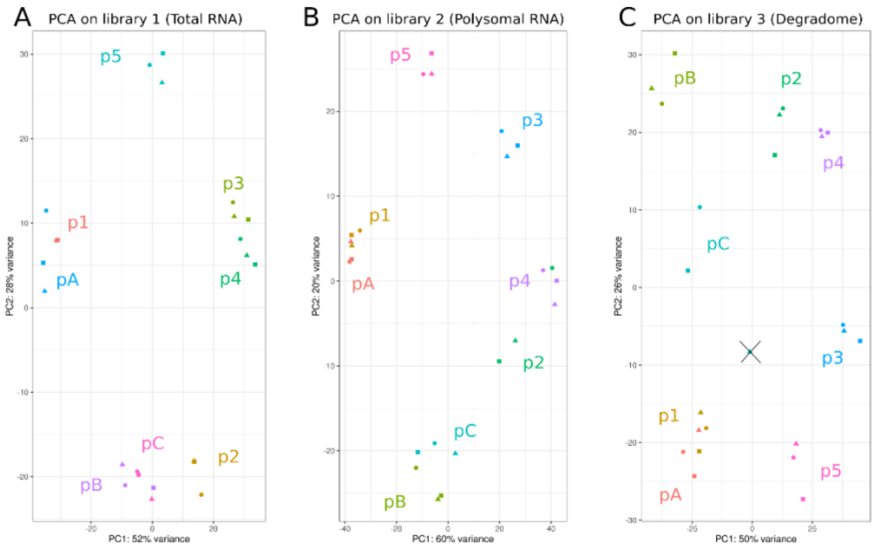


**Supplementary Figure 3. Libraries quality control.** Three two-dimensions PCA were conducted for each library produced (A : library 1, B : library 2 and C : library 3). The two axes can explain 80%, 80 and 76% of the variance for each library respectively. Each replicate is represented by round (R1), triangle (R2), square (R3). In "C", the second replica of pC diverged from the other two and was withdrawn from the study (represented by a cross on the figure).
